# Epigenomic mapping in B-cell acute lymphoblastic leukemia identifies transcriptional regulators and noncoding variants promoting distinct chromatin architectures

**DOI:** 10.1101/2023.02.14.528493

**Authors:** Kelly R. Barnett, Robert J. Mobley, Jonathan D. Diedrich, Brennan P. Bergeron, Kashi Raj Bhattarai, Wenjian Yang, Kristine R. Crews, Christopher S. Manring, Elias Jabbour, Elisabeth Paietta, Mark R. Litzow, Steven M. Kornblau, Wendy Stock, Hiroto Inaba, Sima Jeha, Ching-Hon Pui, Charles G. Mullighan, Mary V. Relling, Jun J. Yang, William E. Evans, Daniel Savic

## Abstract

B-cell lineage acute lymphoblastic leukemia (B-ALL) is comprised of diverse molecular subtypes and while transcriptional and DNA methylation profiling of B-ALL subtypes has been extensively examined, the accompanying chromatin landscape is not well characterized for many subtypes. We therefore mapped chromatin accessibility using ATAC-seq for 10 B-ALL molecular subtypes in primary ALL cells from 154 patients. Comparisons with B-cell progenitors identified candidate B-ALL cell-of-origin and AP-1-associated *cis*-regulatory rewiring in B-ALL. *Cis*-regulatory rewiring promoted B-ALL-specific gene regulatory networks impacting oncogenic signaling pathways that perturb normal B-cell development. We also identified that over 20% of B-ALL accessible chromatin sites exhibit strong subtype enrichment, with transcription factor (TF) footprint profiling identifying candidate TFs that maintain subtype-specific chromatin architectures. Over 9000 inherited genetic variants were further uncovered that contribute to variability in chromatin accessibility among individual patient samples. Overall, our data suggest that distinct chromatin architectures are driven by diverse TFs and inherited genetic variants which promote unique gene regulatory networks that contribute to transcriptional differences among B-ALL subtypes.

**HIGHLIGHTS:**

- Pro-B progenitor cells as the most common cell-of-origin for B-ALL
- AP-1 TF-associated *cis*-regulatory rewiring in B-ALL
- Subtype-specific accessible chromatin signatures representing 20% of all B-ALL sites
- Role for distinct TFs in promoting subtype-specific chromatin architectures
- Thousands of inherited genetic variants identified impacting chromatin state

## INTRODUCTION

Acute lymphoblastic leukemia (ALL) is derived from B- and T-cell lineage precursor cells and is the most common childhood cancer ^1^. A majority of acute lymphoblastic leukemias are derived from B-cell lineages (B-ALL) that are comprised of distinct molecular subtypes characterized by unique chromosomal lesions, including aneuploidy, translocations, gene fusions, point mutations and other chromosomal rearrangements that drive leukemogenesis ^2^. Numerous studies have identified extensive heterogeneity in transcriptomes ^3,4^ and DNA methylomes ^5,6^ among B-ALL subtypes in large patient cohorts, but there is limited understanding of chromatin landscapes. Here we provide an extensive survey of accessible chromatin state and *cis*-regulatory element activity in primary B-ALL cells from over 150 patients across the United States.

Chromatin accessibility or open chromatin is a hallmark of active *cis*-regulatory elements that control spatial and temporal gene expression ^7^. Because ALL typically involves mutations (*PAX5-* altered), complex rearrangements (*DUX4-*rearranged, *PAX5-*altered, *ZNF384-*rearranged, etc.) and/or oncogenic gene fusions (*ETV6∷RUNX1*, *TCF3∷PBX1*, *KMT2A-*rearranged, etc.) of transcription factor (TF) genes as well as disruptions of *cis*-regulatory elements^8^, chromatin accessibility maps can provide valuable information to better understand the leukemogenic process. Accessible chromatin sites can be mapped using transposases by performing assay for transposase-accessible chromatin with high-throughput sequencing (ATAC-seq) ^9,10^. Although DNase treatment has also been used ^11^, one key advantage of ATAC-seq is the low sample input requirements compared to DNase-based assays. This makes ATAC-seq an attractive assay for mapping open chromatin in primary cells from patients wherein sample availability is limited. Additionally, chromatin accessibility allows for identification of bound TFs through an examination of TF footprints which are defined by a depletion in DNA transposition ^12^ or DNase ^13^ cleavage events within regions of accessible chromatin signal. As a result, the underlying TF-binding gene regulatory networks that promote chromatin accessibility and differential gene expression can be predicted.

Previous large-scale studies of chromatin accessibility in primary cells have predominantly focused on distinct cell types ^10,14^ or distinct tumor types and locations ^15,16^. Therefore, large-scale analyses aimed to better understand chromatin state in a single heterogeneous malignancy are currently lacking. To address this knowledge gap, we mapped chromatin accessibility in fresh primary ALL cells from 154 patients across 10 molecular subtypes of B-ALL (*BCR∷ABL1*, *DUX4-*rearranged, *ETV6∷RUNX1*, high hyperdiploid, low hypodiploid, *KMT2A-*rearranged, *BCR∷ABL1*-like (Ph-like), *PAX5-*altered, *TCF3∷PBX1*, *ZNF384-*rearranged) and B-other patient samples. Notably, these subtypes span the entire spectrum of clinical prognoses, including patients with excellent (*DUX4-* rearranged, *ETV6∷RUNX1*, high hyperdiploid), good (*TCF3∷PBX1*), intermediate (*ZNF384-*rearranged, *PAX5-*altered) and poor (*BCR∷ABL1*, low hypodiploid, *KMT2A-*rearranged and Ph-like) prognosis. We also mapped histone H3 lysine 27 acetylation (H3K27ac) enrichment using ChIP-seq in a subset of these patient samples to additionally infer functional activity.

Using ATAC-seq chromatin accessibility and histone profiling in primary ALL cells, we mapped *cis*-regulatory element activity in B-ALL. In complement to chromatin accessibility profiling, we identified thousands of chromatin loops targeting promoters in multiple B-ALL cell lines to better inform linkages of *cis*-regulatory elements to cognate genes. We coupled these maps to transcription factor (TF) footprints at accessible chromatin sites to identify key TFs and gene regulatory networks across B-ALL samples and within distinct B-ALL subtypes. Our results identified extensive chromatin reprogramming between B-cell progenitors and B-ALL, as well as extensive heterogeneity in accessible chromatin landscapes among B-ALL subtypes. Specifically, we uncovered a focused subset of over 42,000 B-ALL open chromatin sites exhibiting extensive subtype-enrichment and subtype-enriched TF binding events. Notably, these sites can predict and classify B-ALL samples with 86% cross-validation accuracy. We additionally explored the impact of inherited genetic variation on chromatin state and delineated over 9000 ATAC-seq chromatin accessibility quantitative trait loci (ATAC-QTLs) in B-ALL cells, including a subset that alter neighboring gene expression. Using the largest accessible chromatin accessibility dataset for B-ALL to date, our data collectively support substantial subtype-specificity in chromatin accessibility that is driven in part by distinct TFs, as well as pronounced inter-individual heterogeneity in chromatin state through inherited genetic variants. Our work further supports the role of these distinct chromatin architectures in establishing unique gene regulatory networks that impact gene expression and B-ALL cell biology.

## RESULTS

### Chromatin accessibility profiles of B-ALL patient samples spanning multiple subtypes

ATAC-seq using the Fast-ATAC ^10^ method was performed on recently-harvested primary ALL cells from 154 patients spanning 10 B-ALL molecular subtypes (*BCR∷ABL1*, *DUX4-*rearranged, *ETV6∷RUNX1*, high hyperdiploid, low hypodiploid, *KMT2A-*rearranged, Ph-like, *PAX5-*altered, *TCF3∷PBX1*, *ZNF384-* rearranged) and B-other samples (**Table S1**) from diverse medical centers, research groups and clinical trials networks across the United States (see **Methods**). To identify high-confidence sites, we identified ATAC-seq peak summits using subtype merged data and selected only loci reproducible among unmerged individual patients. Using this approach we identified 110,468 accessible chromatin fsites, on average, in each B-ALL subtype (range= 71,797–142,498), with 217,240 merged sites identified in total representing the final genomic regions of interest (**Figure 1A, Table S2**).

**FIGURE 1:**
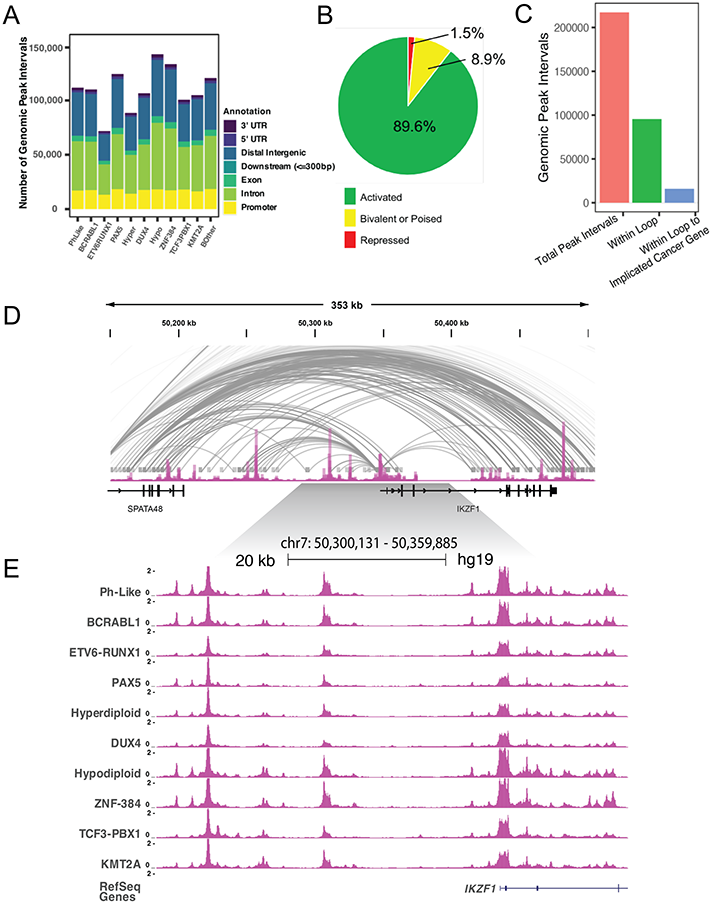
Chromatin accessibility landscapes in B-ALL. **(A)** Number and genomic location of accessible chromatin sites for 10 B-ALL subtypes and B-other samples is provided. **(B)** Percentage of B-ALL accessible chromatin sties that maps to H3K4me1 and/or H3K27ac active histone marks (Active; green), H3K27me3 and H3K4me1 and/or H3K27ac bivalent or poised histone marks (Bivalent or Poised; yellow) and H3K27me3 only repressed histone marks (Repressed; red). **(C)** B-ALL cell line chromatin loops detected using promoter capture Hi-C at B-ALL accessible chromatin sites. The total number of B-ALL accessible chromatin sites, number of B-ALL accessible chromatin sites within loops and the total number of accessible chromatin sites with a loop to a gene implicated in cancer is shown. **(D)** UCSC genome browser ATAC-seq signal track of average B-ALL chromatin accessibility and promoter capture Hi-C loops across the *IKZF1* gene locus. **(E)** UCSC genome browser ATAC-seq signal tracks of 10 merged B-ALL subtypes with known molecular drivers across the *IKZF1* gene locus.

Using H3K27ac ChIP-seq data generated from a subset of 11 B-ALL patient samples, as well as primary B-ALL cell H3K27ac, H3K4me1 and H3K27me3 ChIP-seq data from the Blueprint Epigenome Consortium (https://www.blueprint-epigenome.eu/), we determined that nearly all open chromatin sites mapped to regions containing only active histone marks (H3K27ac and/or H3K4me1, 89.6%; H3K27ac= 3.3%, H3K4me1=34% and H3K4me1+H3K27ac=52.3%) or regions with bivalent marks suggesting a poised chromatin state (H3K27ac and/or H3K4me1 and H3K27me3, 8.9%), compared to only 1.5% of ATAC-seq sites that mapped to regions solely harboring repressive chromatin (H3K27me3; **Figure 1B**). Because these histone modifications are typically found at transcriptional enhancers and promoters^17–20^, these findings suggest that these accessible chromatin regions are B-ALL *cis*-regulatory elements implicated in gene regulation.

In most cases, these candidate *cis*-regulatory elements map within intergenic or intragenic loci with unclear gene targets. Therefore, to better inform gene connectivity we produced chromatin looping data using promoter capture Hi-C ^21^ across seven B-ALL cell lines (697, BALL1, Nalm6, REH, RS411, SEM and SUPB15) to complement B-ALL patient chromatin accessibility profiles. Collectively, across the B-ALL cell lines we detected approximately 400,000 chromatin loops, with approximately 50% of the 217,240 chromatin accessible regions of interest intersecting with a promoter loop, including 15,929 chromatin accessible sites that looped to a cancer implicated gene set (**Figure 1C**) ^22,23^. In many instances, large domains of extensive chromatin looping are present, which with chromatin accessibility and active histone marks emphasize the gene regulatory networks present across B-ALL patient samples (e.g., **Figures 1D** and **1E**).

### Chromatin accessibility identifies Pro-B cell-of-origin for most B-ALL patient samples

To better understand chromatin remodeling during leukemogenesis we sought a comparison of chromatin accessibility between B-ALL and B-cell progenitors. Moreover, although it is widely accepted that the B-ALL cell-of-origin is a B-cell precursor, exactly which precursor is not always clear, particularly at the chromatin accessibility level ^24^. To resolve this uncertainty, we examined publicly available ATAC-seq data from several human B-cell progenitors^10,25^ (**Figure 2A**). When comparing chromatin accessibility signal between B-cell progenitor groups, we identified a set of approximately 42,344 genomic loci which demonstrate a chromatin accessibility enrichment or depletion trend for a B-cell progenitor (**Figure 2B, Table S3**). We refer to these chromatin loci as B-progenitor identity loci due their distinct patterning across B-progenitor differentiation and are likely representations of stage-specific gene regulatory programs.

**FIGURE 2:**
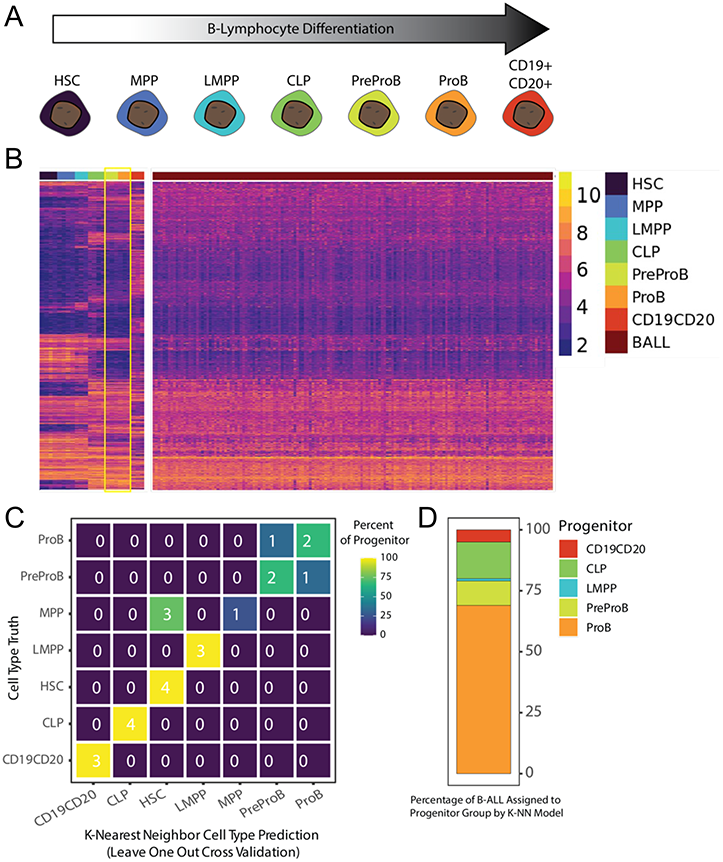
B-ALL cell type-of-origin defined by chromatin accessibility. **(A)** Differentiation timeline of B-cell progenitors from least differentiated to most differentiated. HSC= hematopoietic stem cell, MPP= multipotent progenitor cell, LMPP= lymphoid-primed multipotent progenitor cell, CLP= common lymphoid progenitor cell, PreProB= prePro-B cell, ProB= Pro-B cell and CD19+,CD20+= B cell. **(B)** Heatmap of B-cell progenitor or B-ALL patient sample variance stabilized ATAC-seq signal across B-cell progenitor-defining chromatin loci. B-cell progenitor groups most similar to B-ALL patient samples (preProB and ProB) are outlined in yellow. **(C)** Confusion matrix showing number (listed) and percentage (color-coded) of B-cell progenitor truths and predictions for leave-one-out cross validation of a K-nearest neighbor classifier model. **(D)** Distribution of B-cell progenitor classification across B-ALL patient samples using a K-nearest neighbor classifier model trained with B-cell progenitor data.

Next, we examined patient B-ALL cell chromatin accessibility across these B-progenitor identity loci. When plotting chromatin accessibility signal as a heatmap comparing B-cell progenitors and B-ALL patient samples, a high degree of similarity was observed with prePro-B cells and Pro-B cells (**Figure 2B**). Further, when applying the K-nearest neighbor classification model previously trained on B-progenitor identity loci the majority of B-ALL samples classified as either prePro-B or Pro-B (**Figures 2C** and **2D**). However, prePro-B cells have been reported to be an extremely rare population beyond embryonic and fetal development ^25^. Overall, Pro-B cells demonstrate the most similarity to B-ALL cells at the chromatin accessibility level when focusing specifically on B-cell precursor defining loci, emphasizing this precursor B-cell as a common cell-of-origin for B-ALL.

### Extensive differences in chromatin accessibility between B-ALL and Pro-B cells

To better understand chromatin remodeling during leukemogenesis we next compared accessible chromatin sites between B-ALL and Pro-B cells (n=3) and uncovered 42,661 differentially accessible chromatin sites (DAS) exhibiting lesser or greater accessibility in B-ALL samples (**Figures 3A** and **3B**; **Figure S1** and **Table S4**). Ontology analysis focusing strictly on DAS with higher chromatin accessibility in B-ALL indicated an enrichment for sites associated with genes involved with toll-like receptor signaling, interleukin production, metabolism (acetyl-CoA production) and cell proliferation (**Figure 3C**). Enriched ontology terms were frequently present at multiple fold change thresholds of input B-ALL DAS (**Table S5)**.

**FIGURE 3:**
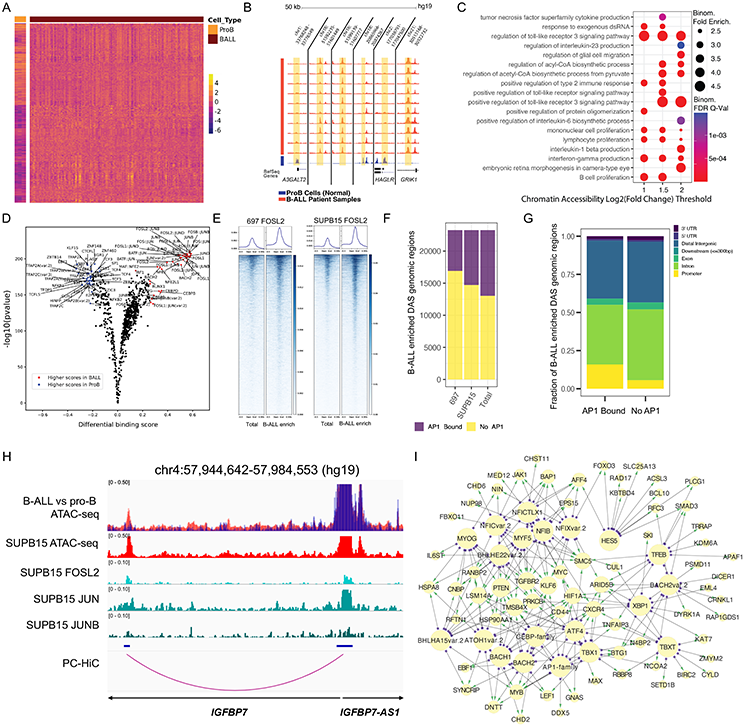
Mapping differential accessibility between B-ALL and Pro-B cells. **(A)** Heatmap of Pro-B cell or B-ALL patient sample variance stabilized ATAC-seq signal as z-score across Pro-B cell and B-ALL enriched DAS. DAS within heatmap are > 1 or < −1 log_2_-adjusted fold change. **(B)** ATAC-seq signal track examples of Pro-B-cell-enriched DAS and B-ALL-enriched DAS on the UCSC genome browser. Flanking genomic regions are included for context. **(C)** Gene ontology analysis of DAS with higher accessibility in B-ALL (B-ALL-enriched) at various log_2_-adjusted fold change thresholds. All terms were significant using both binomial and hypergeometric statistical tests. **(D)** Differential transcription factor footprinting between Pro-B cells and B-ALL patient samples across 217,240 B-ALL genomic regions of interest. **(E)** FOSL2 CUT&RUN enrichment heatmaps at all B-ALL accessible chromatin sites and B-ALL enriched DAS (B-ALL enrich) in SUPB15 (left) and 697 (right) cells. **(F)** Number of B-ALL enriched DAS overlapping AP-1 TF occupancy (FOSL2, JUN and/or JUNB) in 697 (left) SUPB15 (middle) and both B-ALL cell lines (right). Number of overlapping sites are shown in purple while non-overlapping sites are shown in yellow. **(G)** Genome annotation of B-ALL enriched DAS with AP-1 TF occupancy (left) or that are devoid of AP-1 TF occupancy (right). **(H)** IGV genome browser image showing a B-ALL enriched DAS that maps to accessible chromatin and sites of AP-1 TF occupancy in SUPB15 cells. Promoter capture Hi-C (PC-HiC) looping between the distal AP-1 occupied sites and the *IGFBP7* gene promoter is shown. B-ALL (red) and pro-B (blue) cell ATAC-seq tracks are overlaid in the top panel. Signal tracks for FOSL2, JUN and JUNB in SUBP15 cells are shown. **(I)** Transcription factor and target gene network of DAS with higher accessibility in B-ALL (B-ALL-enriched). Network is subset for top transcription factor footprints across DAS ranked by the top mean log_2_-adjusted fold change transcription factor footprint signal. Target genes are subset for a cancer implicated gene set ranked by the top expressed genes. Network connections are colored as transcription factors (purple blocks) to target gene (green arrow heads) pairs. Select expansive and highly similar transcription factor motif families are grouped (AP-1 and CEBP; AP1-family and CEBP-family).

In addition to profiling differential chromatin accessibility, global transcription factor (TF) binding was also compared between B-ALL and Pro-B cells. To identify differential TF binding, we performed genome-wide TF footprint profiling ^12^ using 810 TF motifs comparing B-ALL patient samples and normal Pro-B cell samples across all B-ALL genomic regions of interest (217,240 regions). Differential binding scores indicated the AP-1 family of TFs (e.g., FOS, JUN) as the most prominent TFs with higher binding in B-ALL patient samples compared to normal Pro-B cells (**Figure 3D**). In contrast, prominent TFs with higher binding in Pro-B cells were TFs such as TFAP2A, KLF15, CTCFL, ZBTB14 and EBF1.

To further demonstrate AP-1 TF occupancy in B-ALL accessible chromatin sites we performed CUT&RUN for FOSL2, JUN and JUNB in 697 and SUB15 human B-ALL cell lines (**Figure 3E**; **Figure S2**). Intersections with B-ALL accessible chromatin sites from primary cells identified that 27% of these sites were occupied by an AP-1 TF in B-ALL cell lines. Strikingly, our results further uncovered that 45% of DAS with higher chromatin accessibility in B-ALL (i.e., B-ALL enriched DAS) also exhibit AP-1 TF occupancy (**Figure 3F**), thereby supporting AP-1-associated *cis*-regulatory rewiring in B-ALL. We determined that even though most AP-1 occupied B-ALL enriched DAS localized to promoter-distal regions of the human genome (77%), there is a 2.7-fold enrichment for AP-1 occupancy at B-ALL enriched promoters compared to B-ALL enriched DAS devoid of AP-1 occupancy (**Figure 3G**; 16% vs 6%). Further integration of AP-1 occupied B-ALL enriched DAS with promoter capture Hi-C in B-ALL cell lines identified target genes that were enriched for cell cycle, autophagy and apoptotic signaling pathways (**Table S6**; example in **Figure 3H**).

As an extension of our TF footprinting data we also integrated B-ALL cell line promoter capture Hi-C using the ABC enhancer algorithm to refine identification of TF-target gene relationships across top TFs and a cancer implicated gene set ^26^. Specifically, we focused on top TF footprints within B-ALL enriched DAS and the cancer implicated gene targets of these DAS predicted by the ABC enhancer algorithm. Concordant with global TF footprint and AP-1 TF occupancy analyses we identified the AP-1 family as top TFs in this network. We also identified other top TFs from TF footprinting such as CEBP family TFs and BACH2 (**Figure 3I**). Other prominent top TFs include NFIC, XBP1, TBX1 and numerous basic helix-loop-helix (bHLH) class TFs (e.g., MYOG, MYF5 and HES5). Top expressed cancer implicated gene targets for each TF converged on notable genes involved in cell signaling (*TGFBR2, CXCR4*), histone mark modification (*ARID5B*), transcriptional regulation (*MYC*, *KLF6*, *HIF1A*) and the PI3K-AKT pathway (*PTEN*) (**Figure 3I**). Collectively, these results highlight a rewiring of signaling pathways and TF binding networks that facilitate the proliferative potential of B-ALL samples compared to Pro-B cells.

### Identification of subtype-enriched chromatin architecture

To better understand chromatin accessibility within B-ALL, inter-subtype analyses were performed to identify DAS exhibiting subtype-enriched signal (i.e., henceforth referred to as subtype-enriched DAS) in 10 B-ALL molecular subtypes harboring known molecular drivers (*BCR∷ABL1*, *DUX4-*rearranged, *ETV6∷RUNX1*, high hyperdiploid, low hypodiploid, *KMT2A-*rearranged, Ph-like, *PAX5-*altered, *TCF3∷PBX1* and *ZNF384-*rearranged; **Figures 4A** and **4C**). For this analysis, we compared a single B-ALL subtype cohort with all other B-ALL cell samples not belonging to that subtype in pairwise fashion covering all subtypes. This approach was utilized to emphasize high degrees of subtype enrichment compared to the full spectrum of chromatin accessibility variability in the remaining sample cohort. We identified between 307 and 10,639 DAS in each B-ALL subtype, with a total of 42,457 subtype-enriched DAS identified across all 10 B-ALL subtypes (log_2_ fold change > or < 1, FDR<0.05; **Figure 4B**, **Table S7**). We annotated subtype-enriched DAS on a subtype basis and determined that a majority of subtype-enriched DAS in each B-ALL subtype (87%, range=80%-90%) localized to promoter-distal regions of the genome (intronic and distal intergenic; **Figure 4D**), and 43%, on average (range=39%-49%), localized to distal intergenic regions, thereby emphasizing the importance of non-genic loci in defining B-ALL chromatin heterogeneity.

**FIGURE 4:**
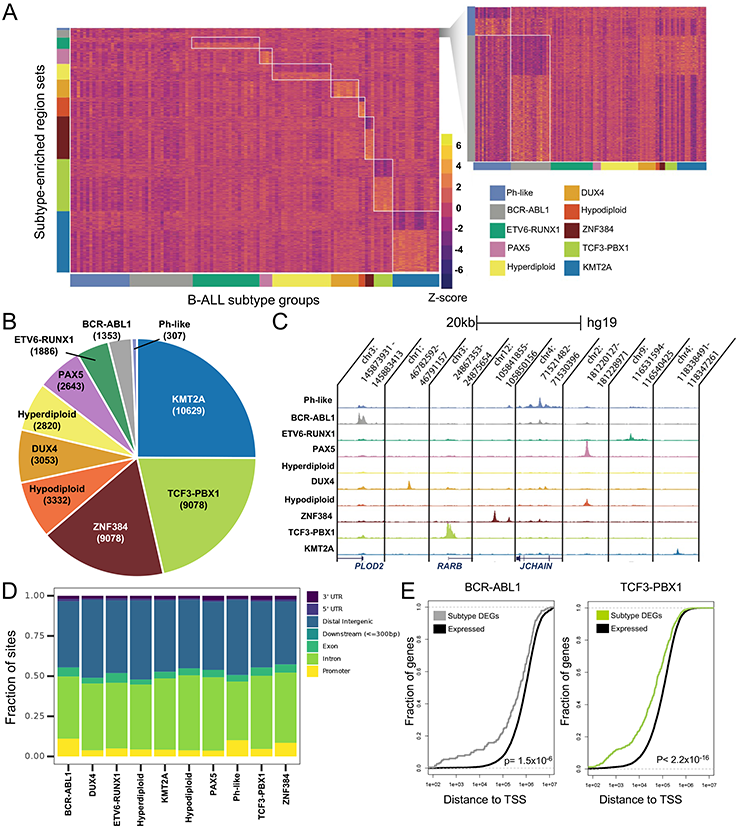
Mapping differential accessibility among B-ALL molecular subtypes. **(A)** Heatmap of variance stabilized ATAC-seq signal as z-score across subtype-enriched DAS. Enrichment patterns for each subtype DAS set are shown on vertical axis and are grouped by B-ALL subtype patient sample on the horizontal axis. Ph-like and BCR-ABL subtype-enriched DAS are expanded at the right for clarity. **(B)** Pie chart shows the number and percentage of subtype-enriched DAS identified. **(C)** ATAC-seq signal track examples of subtype-enriched DAS on the UCSC genome browser. **(D)** Genomic annotations of subtype-enriched DAS for each B-ALL subtype is provided. The fraction of sites harboring different annotations is plotted. **(E)** Cumulative distribution function for *BCR∷ABL1* and *ZNF384-*rearranged ALL comparing the fraction (y-axis) of subtype up-regulated genes (Subtype DEGs; gray or light green) and all expressed subtype genes (Expressed; black) at different distance cutoffs from subtype-enriched DAS and their transcription start sites (x-axis). Kolmogorov-Smirnov (K-S) p-values are provided.

To further evaluate subtype-enriched DAS, we determined if they displayed enrichment patterns that were consistent with five established human B-ALL cell lines (697= *TCF3∷PBX1*, Nalm6= *DUX4-* rearranged, REH= *ETV6∷RUNX1*, SEM= *KMT2A-*rearranged and SUPB15= *BCR∷ABL1*). Concordant with DAS in patient samples, subtype-enriched DAS exhibited the strongest (BCR-ABL, *DUX4-* rearranged, *ETV6∷RUNX1*, *KMT2A-*rearranged) or second strongest (*TCF3∷PBX1*) accessibility in the concordant cell line that was representative of that subtype (**Figure S3**). These data suggest that B-ALL cell lines exhibit chromatin accessibility that is largely consistent with primary B-ALL cell sample from the corresponding subtype.

To further determine functional effects on gene expression, we integrated subtype-enriched DAS with DEGs uniquely up-regulated (log_2_ fold change >1, FDR<0.05) in each of the 10 B-ALL molecular subtypes to determine if they were enriched near DEGs. We identified a statistically significant enrichment of subtype-enriched DAS near up-regulated DEGs in 9 of 10 subtypes compared to total expressed genes in the corresponding subtype (Kolmogorov-Smirnov test p< 0.05; **Figure 4E**, **Figure S4**) and uncovered a strong statistical trend in Ph-like B-ALL (Kolmogorov-Smirnov test p= 0.06; **Figure S4**). Consequently, these data support the role of subtype-enriched DAS in gene regulation and gene activation and further suggest that differences in chromatin accessibility contribute to transcriptomic differences among B-ALL subtypes ^3,4^. Collectively, these results highlight extensive open chromatin heterogeneity among B-ALL molecular subtypes.

### Mapping transcription factor drivers and gene regulatory networks in B-ALL subtypes

We performed TF footprint profiling using merged ATAC-seq signal from 10 B-ALL subtypes with known molecular drivers to identify subtype-enriched TF drivers. TF footprint profiling ^12^ identified between 4,303,155 and 5,441,937 bound motifs in each B-ALL subtype, with 49,402,067 TF footprints at 815,992 unique genomic loci identified across all subtypes. Using these data, we next identified key TF footprints that were enriched in each subtype (i.e., subtype-enriched TF footprints) by calculating differential footprint scores between every subtype-subtype pair for each TF motif. The top median differential motif scores for each subtype were selected as subtype-enriched TF footprints. This approach was utilized to emphasize differential TF footprint motifs that were consistent and distinct for each subtype rather than repetitive global trends (**Figure 5A**). Notably, subtype-enriched TF footprints were identified for recognized TF drivers such as DUX4 in *DUX4-*rearranged ALL and ZNF384 in *ZNF384-*rearranged ALL. We also identified HOX family TFs (HOXA9, HOXB9, HOXC9 and HOXD9) in *KMT2A-*rearranged ALL, GATA family TFs (GATA2, GATA3, GATA4, GATA5 and GATA6) in *ZNF384-* rearranged ALL and nuclear receptor family TFs in *PAX5-*altered ALL (ESR1, ESR2, ESRRA, NR2F6, NR2F1, RARA and THRB) that all had strong subtype-enriched TF footprints.

**FIGURE 5:**
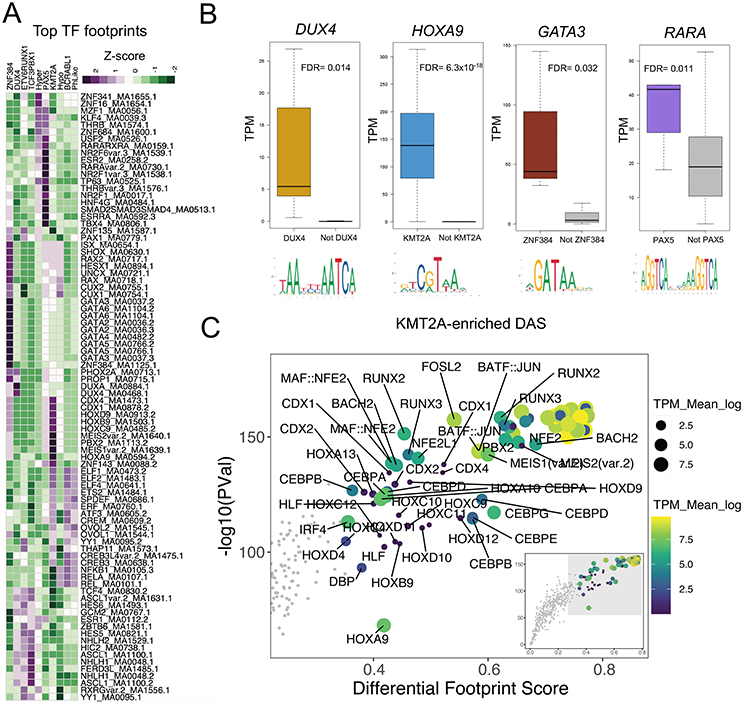
TF footprinting and gene regulatory networks identify key TF drivers in B-ALL subtypes. **(A)** Heatmap list of the topmost consistently differential TF footprints between all pairwise subtype-subtype comparisons (y-axis; labeled to the right of the heatmap as TF motif identifiers) enriched in 10 B-ALL subtypes (x-axis; labeled on top of heatmap as z-score of differential TF footprint signal output by TOBIAS). **(B)** RNA-seq transcripts per million (TPM) expression of key TFs with subtype-enriched footprints that are also up-regulated in the corresponding subtype (colored) versus all other subtypes (gray). DESeq2 differentially expressed gene FDR significance values are provided. **(C)** Top TF footprints at *KMT2A-*enriched DAS are shown. Differential footprint score between B-ALL and Pro-B cells is provided on the x-axis and TF footprint significance is provided on the y-axis. Transcripts per million (TPM) transcript abundance of associated TF transcript is shown as both color and size of points.

Because DNA consensus motifs can be highly redundant within TF families, we integrated subtype-enriched TF footprints with DEGs uniquely up-regulated in each subtype to identify candidate TFs from these TF families that are up-regulated in the corresponding B-ALL subtype. This analysis identified *HOXA9* and *HOXC9*, *RARA* and *GATA3* as up-regulated genes in *KMT2A-*rearranged, *PAX5-* altered and *ZNF384-*rearranged subtypes, respectively (**Figure 5B**, **Figure S5**). In addition, *DUX4* (*DUX4-*rearranged) and *MEIS1* (*KMT2A-*rearranged) were also identified as up-regulated TF genes with subtype-enriched TF footprints (**Figure S5**).

To determine if these up-regulated TFs promote unique chromatin accessibility landscapes among B-ALL subtypes, we also performed TF footprinting on subtype-enriched DAS by comparing differential footprint scores at subtype-enriched DAS between each B-ALL subtype and Pro-B cells (**Figure 5C**, **Figure S6**). Notably, these data supported a role of DUX4 in *DUX4-*rearranged ALL, ZNF384 and GATA3 in *ZNF384-*rearranged ALL, and HOXA9 and MEIS1 in *KMT2A-*rearranged ALL in the generation of subtype-specific chromatin landscapes (**Figure 5C**, **Figure S6**).

### Predictive potential of B-ALL subtype-enriched DAS

We determined how well chromatin accessibility can predict B-ALL subtypes by constructing a stepwise Principal Component Analysis-Linear Discriminant Analysis (PCA-LDA) classification model using the 42,457 subtype-enriched DAS ATAC-seq read count matrix as initial input across 10 B-ALL subtypes harboring known molecular drivers (outlined in **Figure 6A**). Notably, the constructed classification model was tested with leave-one-out cross validation at an accuracy of 86%. The most common failure was incorrect classification of *BCR∷ABL1* and Ph-like subtypes (**Figure 6B**), as has been observed with other ALL classification algorithms ^27^. Taking this into account by grouping *BCR∷ABL1* and Ph-Like subtype samples into a common class yielded a re-calculated cross validation accuracy of 91%. Visualization of B-ALL subtype separations using select dimensions output by the LDA model demonstrates distinct groupings of related subtypes emphasizing classification model performance (**Figure 6C**).

**FIGURE 6:**
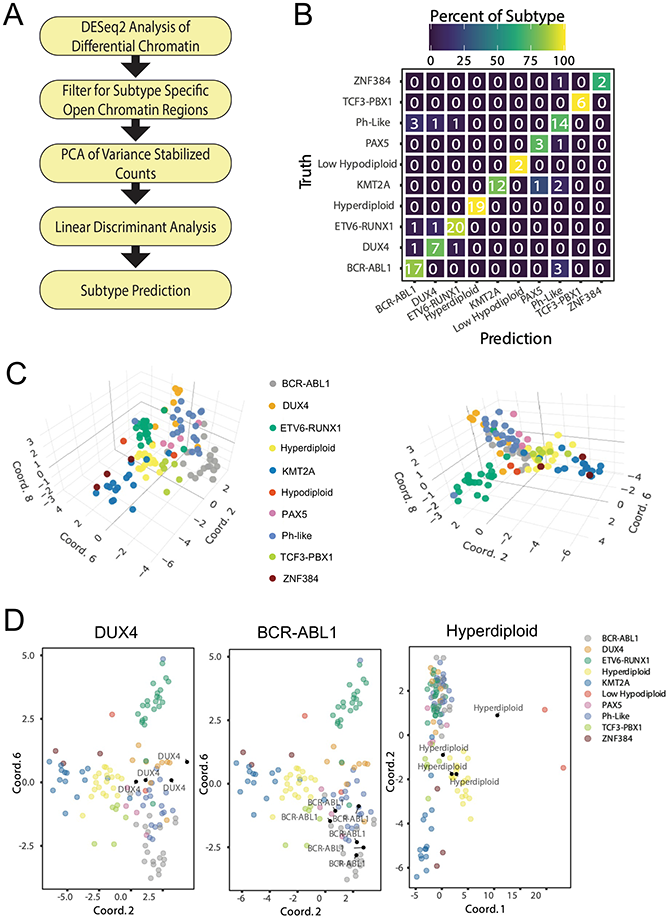
Classification model accurately predicts B-ALL subtypes. **(A)** Flow chart outlines process for PCA-LDA classification of B-ALL subtypes. **(B)** Confusion matrix showing number (listed) and percentage (color-coded) of B-ALL subtype truths and predictions for leave-one-out cross validation. **(C)** Three-dimensional plots showing clustering of B-ALL subtypes utilizing select dimensions from the LDA model. **(D)** B-ALL subtype identification for unknown B-ALL samples (black points). Clustering for unknown samples identified as *DUX4-*rearranged, *BCR∷ABL1* and high hyperdiploid (from left to right) is shown.

As a further application of our classification model, we also applied the algorithm to 26 B-ALL patient samples of unknown molecular B-ALL subtype. Although transcriptomic profiling for B-ALL drivers is not available to fully validate these samples, when processed with the constructed PCA-LDA model and projected onto original LDA dimensions they distinctly cluster with known molecular subtypes supporting reasonable predictions (**Figure 6D**). Collectively, these data support the utility of chromatin structure and subtype-enriched DAS in B-ALL subtype classification.

### Mapping inherited DNA sequence variants that impact chromatin accessibility

To determine how germline variation impacts chromatin accessibility, we identified chromatin accessibility quantitative trait loci using ATAC-seq (ATAC-QTLs) in a subset of 69 patient samples with available SNP genotyping information and allele-specific ATAC-seq read counting using RASQUAL ^28^. In total, 9080 ATAC-QTLs were identified representing both directionalities, with reference or alternative alleles increasing chromatin accessibility (FDR<0.1; **Figure 7A**, **Table S8**). Manual quantification and scaling of allele-specific read counts for select ATAC-QTLs identified with RASQUAL demonstrated a clear concordance and directionality among individual patient samples classified into genotype groups (**Figure 7B**). Visual inspection of merged read counts from patient samples grouped into reference allele homozygote, heterozygote, or alternate allele homozygote for select ATAC-QTLs further supports the high-quality nature of identified ATAC-QTLs (**Figure 7C**). We further determined that 218 ATAC-QTLs where also lead eQTL SNPs when compared to GTEx eQTLs ^29^ from relevant tissues (blood and lymphoblastoid cells), with 85% also concordant for allele overrepresentation directionality (**Figure 7D**; **Table S9**). ATAC-QTLs were also compared with inherited genome-wide association study (GWAS) variants for ALL disease susceptibility which identified rs3824662 (*GATA3*) ^30^ and rs17481869 (2p22.3) ^31^ as ATAC-QTLs that were associated with risk of developing B-ALL. Further supporting the validity of our methodology, rs3824662 was also identified as an ATAC-QTL in ALL PDX samples ^32^, and we functionally validated differential allele-specific activity for rs17481869 in multiple B-ALL cell lines (**Figure S7**).

**FIGURE 7:**
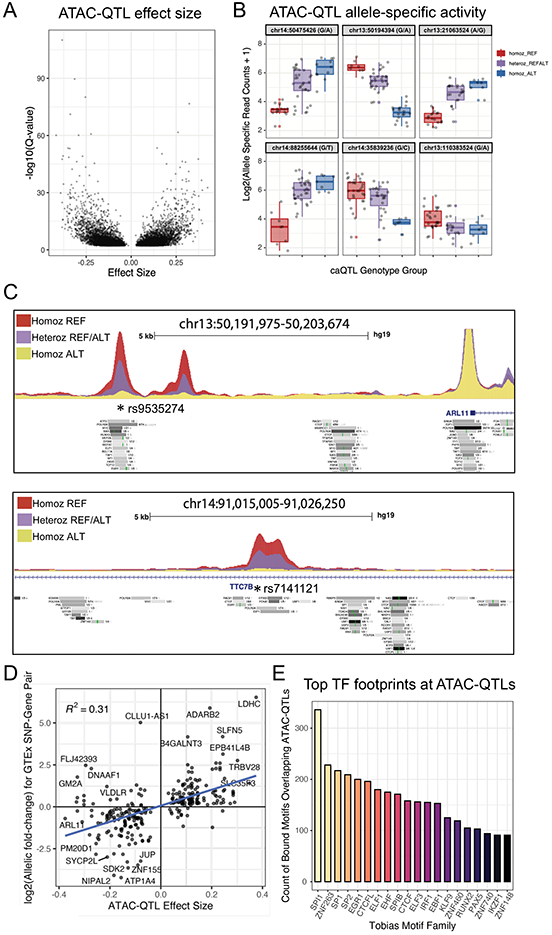
Identification of ATAC-QTLs impacting chromatin accessibility. **(A)** ATAC-QTL effect size (x-axis) and significance (y-axis) is plotted for all significant ATAC-QTLs (FDR<0.1). **(B)** Examples of allele-specific effects on ATAC-seq read count at ATAC-QTLs between samples from the three genotype groups. Homozygous reference allele= homoz_REF, heterozygous= heteroz_REFALT and homozygous alternative allele= homoz_ALT. **(C)** UCSC browser ATAC-seq signal tracks of merged BAM files from patients with distinct genotypes at *ARL11* (top panel) and *TTC7B* (bottom panel) gene loci. ATAC-QTLs are marked by an asterisk. Homozygous reference allele= homoz_REF, heterozygous= heteroz_REFALT and homozygous alternative allele= homoz_ALT. ENCODE ChIP-seq TF binding sites are shown below each ATAC-seq signal track. **(D)** Scatterplot of effect size for SNPs significant as both ATAC-QTLs (x-axis) and GTEx lead eQTL (y-axis). **(E)** Abundance of top TF-bound motifs overlapping ATAC-QTLs. Highly similar TF motifs were grouped into motif families via TOBIAS motif clustering as shown on the x-axis.

To infer the impact of TF binding in control of chromatin accessibility at ATAC-QTLs we overlapped ATAC-QTL loci with TF motifs determined as TF-bound by footprint profiling ^12^. Nearly one-third (28.8%; 2615/9080 ATAC-QTLs) of these ATAC-QTLs overlapped a TF-bound motif footprint across multiple B-ALL subtypes, suggesting that most ATAC-QTLs do not have a clear TF-binding mechanism in how they impact chromatin accessibility. Analysis of bound TF motif footprint prevalence at ATAC-QTLs identified several ETS family TFs (EHF, ELF3, SPI1/PU.1 and SPIB), zinc finger TFs (ZNF263, ZNF460, ZNF740 and ZNF148) and CTCF as the most altered motifs leading to differences in chromatin accessibility between alleles (**Figure 7E**). Notably, we also identified PAX5 and IKZF1, which have known roles in B-cell development and leukemogenesis ^33–36^. Collectively, these data identify inherited DNA sequence variants contributing to chromatin heterogeneity among B-ALL subtypes and indicate specific TFs of interest for further exploration of ATAC-QTLs.

## DISCUSSION

Our study provides the first, large-scale examination of chromatin accessibility in the B-ALL genome across an expansive set of B-ALL subtypes. We further integrated this data with ChIP-seq histone modification enrichment in primary B-ALL cells and three-dimensional chromatin looping data using promoter capture Hi-C in multiple B-ALL cell lines. Our data demonstrate that most regions of chromatin accessibility harbor activating chromatin marks consistent with *cis*-regulatory elements involved in gene regulation, and we further confirmed direct looping to gene promoters for approximately 50% of accessible chromatin sites. However, this does not rule out more transient chromatin looping interactions difficult to detect by current chromatin conformation capture genomic techniques.

Extensive epigenomic reprogramming was uncovered between B-cell progenitors and B-ALL, and cell-of-origin analyses identified Pro-B cells as the most common cell-of-origin. Our comparison of B-ALL and pro-B cell chromatin accessibility suggests epigenomic reprogramming that is, in part, associated with AP-1 TF occupancy. We further identify disruptions to normal B-cell function through the activation of toll-like receptor signaling and interleukin production. Acetyl Co-A synthesis was also identified as an enriched gene ontology term when comparing B-ALL and Pro-B cells. Metabolic alterations in cancer are well known, particularly acetyl-Co-A synthesis alterations which have been previously reported in cancer ^37^. In addition to metabolic alterations, *PTEN*, a known tumor suppressor gene is frequently mutated in a large portion of cancers ^38^. However, in B-ALL the cancer role of PTEN has been reported to be inverted, functioning instead as an oncogene ^39^. Reinforcing this conclusion and further suggesting PTEN as an intriguing target for B-ALL treatment, we also found *PTEN* in our network as a top gene target of B-ALL enriched DAS.

We further examined accessible chromatin landscapes among diverse molecular subtypes of B-ALL. Collectively, we identified 42,457 subtype-enriched DAS which strikingly represent 20% of analyzed accessible chromatin sites across a pan-subtype B-ALL genome. Subtype-enriched DAS were enriched near up-regulated DEG in the corresponding subtype, supporting their role in gene activation. Moreover, comparisons between subtype-enriched DAS and chromatin accessibility data from cell lines identified largely consistent patterns. We further identified candidate TFs that exhibited strong subtype-specificity through TF footprinting analyses and validated some of these findings using transcriptomic data from primary B-ALL cells. Collectively, these analyses highlighted the role of HOXA9 and MEIS1 in *KMT2A-*rearranged ALL, GATA3 in *ZNF384-*rearranged ALL and RARA in *PAX5-*altered B-ALL. We further confirmed the previously reported roles of DUX4 and ZNF384 in *DUX4-*rearranged and *ZNF384-*rearranged ALLs, respectively. Concordant with our findings, previous studies have identified the co-upregulation of HOXA9 and MEIS1 in *KMT2A-*rearranged leukemias and further support that these TFs are key drivers of leukemogenesis ^40–42^. Our identification of numerous HOX TFs with enriched footprints in *KMT2A-*rearranged ALL is also consistent with observations of HOX gene dysregulation in this subtype ^43^. Further supporting our results, ZNF384 fusion proteins in *ZNF384-*rearranged ALL are known to up-regulate *GATA3* expression ^44,45^. Although a direct role for RARA in *PAX5-*altered B-ALL has not been established, previous work has identified *PAX5* as a target gene of the PLZF-RARA fusion protein in acute promyelocytic leukemia ^46^. Moreover, both *RARA* and *PAX5* genes can form fusions with *PML* in acute promyelocytic leukemia ^47^ and ALL ^48^, respectively. While *PAX5-*altered ALL has not been well connected to RARA nuclear receptor signaling, there has been previous work treating IKZF1 mutated BCR-ABL1 ALL with RARA and RXR agonists that suppressed a self-renewal phenotype ^49^. Collectively, these data warrant further investigation of RARA and RXR signaling in *PAX5-*altered ALL.

Supporting the utility of chromatin accessibility in B-ALL classification, subtype-enriched DAS predicted subtypes with 86% accuracy. As a comparison to chromatin accessibility, transcriptional profiling using ALLSorts correctly assigned B-ALL subtypes with 92% accuracy ^27^. However, this RNA-seq dataset included over 1223 transcriptomes from 18 subtypes representing a considerably larger dataset for model development. We therefore suspect that additional chromatin accessibility profiling across more B-ALL subtypes and increased sample sizes will lead to even better subtype prediction that will rival transcriptomic profiling and importantly, incorporate intergenic heterogeneity that can elucidate *cis*-regulatory drivers of B-ALL leukemogenesis.

To identify the role of inherited DNA sequence variation on the B-ALL chromatin landscape, we mapped over 9000 ATAC-QTLs (FDR<0.1). A large subset of ATAC-QTLs mapped to TF footprints and were concordant in allelic biases with GTEx lead eQTLs. Further validating our analysis, we functionally validated a variant (rs17481869; 2p22.3) associated with susceptibility to ALL ^31^. Collectively, this analysis suggests that chromatin accessibility is additionally modified by inherited DNA sequence variation, thereby further contributing to increased chromatin heterogeneity in B-ALL.

Overall, our data support pronounced changes in chromatin accessibility between B-ALL and precursor B-cells, as well as among B-ALL subtypes. Our results further support the role of diverse TFs and inherited genetic variants in modulating and promoting differences in chromatin accessibility among B-ALL subtypes. Ultimately, these diverse chromatin architectures contribute to unique gene regulatory networks and transcriptional programs. Our work therefore provides a valuable resource to the cancer genomics research community and can be further used to better understand biological as well as clinical differences among B-ALL subtypes.

## METHODS

### Patient samples

Patient samples were obtained from: St. Jude Children’s Research Hospital (Memphis, Tennessee), ECOG-ACRIN Cancer Research Group, The Alliance for Clinical Trials in Oncology, MD Anderson Cancer Center (Houston, Texas), Cook Children’s Medical Center (Fort Worth, Texas), Lucile Packard Children’s Hospital (Palo Alto, California), The University of Chicago (Chicago, Illinois), Novant Health Hemby Children’s Hospital (Charlotte, North Carolina) and Children’s Hospital of Michigan (Detroit, Michigan). All patients or their legal guardians provided written informed consent. The use of these samples was approved by the institutional review board at St. Jude Children’s Research Hospital.

### Functional genomic studies

ATAC-seq using the Fast-ATAC^10^ protocol was performed on 10,000 fresh primary ALL cells. H3K27ac ChIP-seq was performed as previously described^50^ on 20 million fresh primary ALL cells. CUT&RUN for FOSL2/Fra2 (Cell signaling; 19967S), JUN (Epicypher; 13-2019) and JUNB (Cell Signaling; 3753S) was performed using the Epicypher Cutana CUT&RUN kit v3.0 (14-1048) according to the manufacturers provided instructions. Next-generation sequencing of ATAC-seq, CUT&RUN, and ChIP-seq was performed at the Hartwell Center for Bioinformatics and Biotechnology at St. Jude Children’s Research Hospital. Transcriptomic and SNP genotyping data from B-ALL patient samples were obtained from St. Jude Children’s Research Hospital. Normal B-cell ATAC-seq ^10,25^ were downloaded from NCBI (GSE122989 and GSE74912). B-ALL cell histone modification ChIP-seq datasets (H3K27ac, H3K4me1 and H3K27me3) were downloaded from the Blueprint Epigenome Consortium (https://www.blueprint-epigenome.eu/). Expression quantitative trait loci (eQTL) data was obtained from previous studies ^51^. Arima promoter capture Hi-C (Arima; A510008, A303010, A302010) was performed on 10 million B-ALL cell lines (697, BALL1, Nalm6, RS411, REH, SEM and SUPB15) according to the manufacturers provided instructions using unspecified proprietary buffers, solutions, enzymes, and reagents. See **Supplemental Methods** for additional details.

### Data analysis

ATAC-seq and ChIP-seq reads were mapped to the hg19 reference genome using bowtie2 ^52^ and peaks were identified using MACS2 ^53^. Regions of interest for ATAC-seq analyses were selected using a reproducible peak summit approach within each subtype cohort with subsequent region merging. DESeq2 ^54^ was employed to identify B-ALL-enriched or subtype-enriched differentially accessible chromatin sites (DAS). Two B-ALL subtype patient samples (IKZF1 N159Y and iAMP21) were included in B-ALL versus Pro-B cell analyses but were excluded from additional studies due to limited sample size. Promoter capture Hi-C libraries from B-ALL cell lines were analyzed at 3-kb resolution using the Arima CHiC pipeline (v1.4, https://github.com/ArimaGenomics/CHiC). Genomic regions representing separate loop ends were compiled to facilitate overlap determinations with B-ALL patient chromatin accessible regions of interest using “bedtools intersect”. Enhancer and target gene prediction for network construction was analyzed with the ABC enhancer algorithm (https://github.com/broadinstitute/ABC-Enhancer-Gene-Prediction). In brief, inputs for the ABC enhancer algorithm included, B-ALL enriched DAS, merged B-ALL patient ATAC-seq, H3K27Ac ChIP-seq, Arima promoter capture Hi-C contact counts with ABC score threshold at 0.04. The Genomic Regions Enrichment of Annotations Tool (GREAT) ^55^ was used to identify candidate target gene sets and ontologies associated with DAS. TOBIAS ^12^ was used to identify TF footprints at accessible chromatin sites. The Principal Component Analysis-Linear Discriminant Analysis (PCA-LDA) subtype classification model was constructed stepwise by first PCA transformation of subtype-enriched ATAC-seq counts, then applying LDA on an optimized number of principal components. RASQUAL ^28^ was used to map chromatin accessibility quantitative trait loci using ATAC-seq (ATAC-QTLs). Significant ATAC-QTLs for each region were identified with a genome-wide computed FDR of 10%. See **Supplemental Methods** for additional details.

## DATA AND CODE AVAILABILITY

Further information and requests for resources should be directed to and will be fulfilled by the lead contact, Daniel Savic.

## ACKNOWLEDGEMENTS

We would like to thank the Hartwell Center at St. Jude for ATAC-seq, ChIP-seq and promoter capture Hi-C library preparation and next-generation sequencing. We would also like to thank Jeremy Hunt and Brandon Smart for technical support. This work was supported by the National Cancer Institute (R01CA234490, P30CA021765, UG1CA232760, UG1CA189859, and U10CA180820), the National Institute of General Medical Studies (P50GM115279) and the American Lebanese Syrian Associated Charities (ALSAC). The content is solely the responsibility of the authors and does not necessarily represent the official views of the National Institutes of Health.

## AUTHOR CONTRIBUTIONS

Conceptualization, K.R. Barnett, D.S.; Methodology, K.R. Barnett, D.S.; Investigation, K.R. Barnett, J.D.D., B.P.B, K.R. Bhattarai; Analysis, K.R. Barnett, D.S.; Data Curation, K.R.B., W.Y.; Patient sample acquisition, K.R.C., C.S.M., E.J., E.P., M.R.L., S.M.K., W.S., H.I., S.J., C.H.P., C.G.M., M.V.R., W.E.E., J.J.Y.; Writing – Original Draft, K.R. Barnett, D.S.; Writing – Review & Editing, K.R. Barnett, J.D.D., B.P.B, K.R. Bhattarai, W.Y., K.R.C., C.S.M., E.J., E.P., M.R.L., S.M.K., W.S., H.I., S.J., C.H.P., C.G.M., M.V.R., W.E.E., J.J.Y, D.S.

