## Supplemental Figures for "Epigenomic mapping in B-cell acute lymphoblastic leukemia identifies transcriptional regulators and noncoding variants promoting distinct chromatin architectures"

**Chromatin accessibility landscapes of B-cell acute lymphoblastic leukemia subtypes identifies epigenomic heterogeneity**

Kelly R. Barnett, PhD^1,2^, Robert J. Mobley, PhD^1,2^, Jonathan D. Diedrich, PhD^1,2^, Brennan P. Bergeron, PhD^1,2,3^, Kashi Raj Bhattarai, PhD^1,2^, Wenjian Yang, PhD^1,2^, Kristine R. Crews, PharmD^1,2^, Christopher S. Manring, MBA^4^, Elias Jabbour, MD^5^, Elisabeth Paietta, PhD^6^, Mark R. Litzow, MD^7^, Steven M. Kornblau, MD^5^, Wendy Stock, MD^8^, Hiroto Inaba, MD, PhD^1,9^, Sima Jeha, MD^1,9^, Ching-Hon Pui, MD^1,9^, Charles G. Mullighan, MBBS (Hons), MSc, MD^1,10^, Mary V. Relling, PharmD^1,2^, Jun J. Yang, PhD^1,2,3,11^, William E. Evans, PharmD^1,2^ and Daniel Savic, PhD^1,2,3,11,*^

^1^ Hematological Malignancies Program, St. Jude Children’s Research Hospital, Memphis, TN 38105, USA.

^2^ Department of Pharmacy and Pharmaceutical Sciences, St. Jude Children’s Research Hospital, Memphis, TN 38105, USA.

^3^ Graduate School of Biomedical Sciences, St. Jude Children’s Research Hospital, Memphis, TN 38105, USA.

^4^ Alliance Hematologic Malignancy Biorepository; Clara D. Bloomfield Center for Leukemia Outcomes Research, Columbus, OH 43210, USA

^5^ Department of Leukemia, The University of Texas M. D. Anderson Cancer Center, Houston, TX, USA

^6^ Department of Oncology, Montefiore Medical Center, Bronx, NY 10467, USA.

^7^ Division of Hematology, Department of Medicine, Mayo Clinic, Rochester, MN 55905, USA.

^8^ University of Chicago Comprehensive Cancer Center, Chicago, IL 60637, USA.

^9^ Department of Oncology, St. Jude Children’s Research Hospital, Memphis, TN 38105, USA.

^10^ Department of Pathology, St. Jude Children’s Research Hospital, Memphis, TN 38105, USA.

^11^ Integrated Biomedical Sciences Program, University of Tennessee Health Science Center, Memphis, TN 38105, USA.

*Corresponding author:	Daniel Savic, PhD

Division of Pharmaceutical Sciences

Department of Pharmacy and Pharmaceutical Sciences

St. Jude Children’s Research Hospital

262 Danny Thomas Place

Memphis, TN, 38105

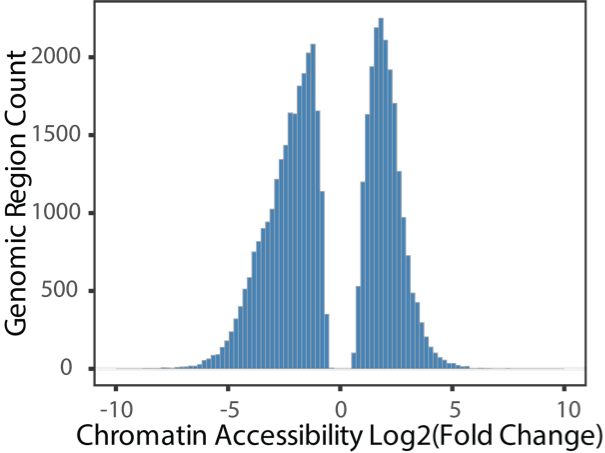

**Figure S1.** Histogram of log_2_-adjusted fold change in ATAC-seq signal at significant DAS between Pro-B cells and B-ALL patient samples.

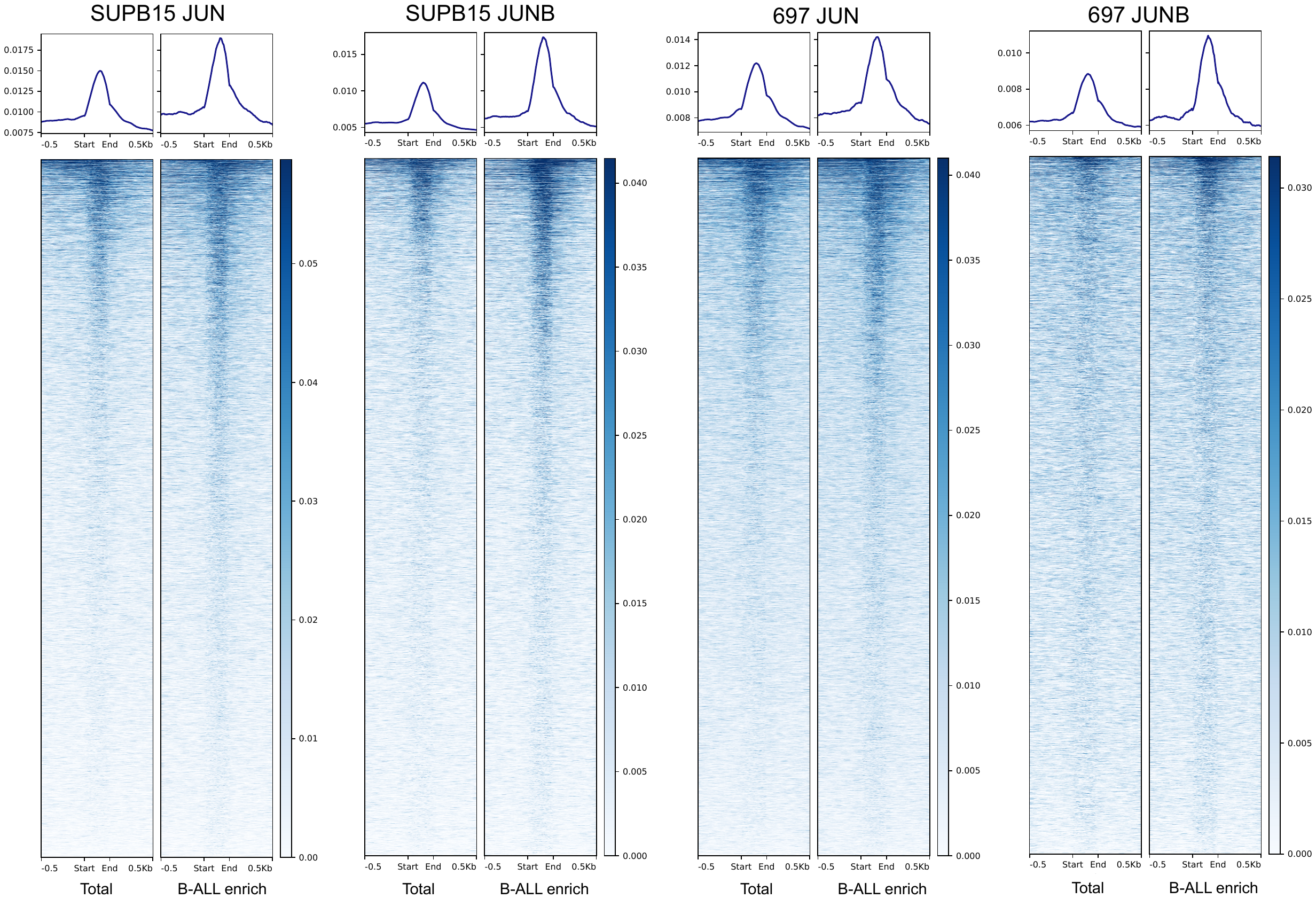

**Figure S2.** JUN and JUNB CUT and RUN enrichment heatmap at all B-ALL accessible chromatin sites and B-ALL enriched DAS (B-ALL enrich) in SUPB15 (left) and 697 (right) cells is shown.

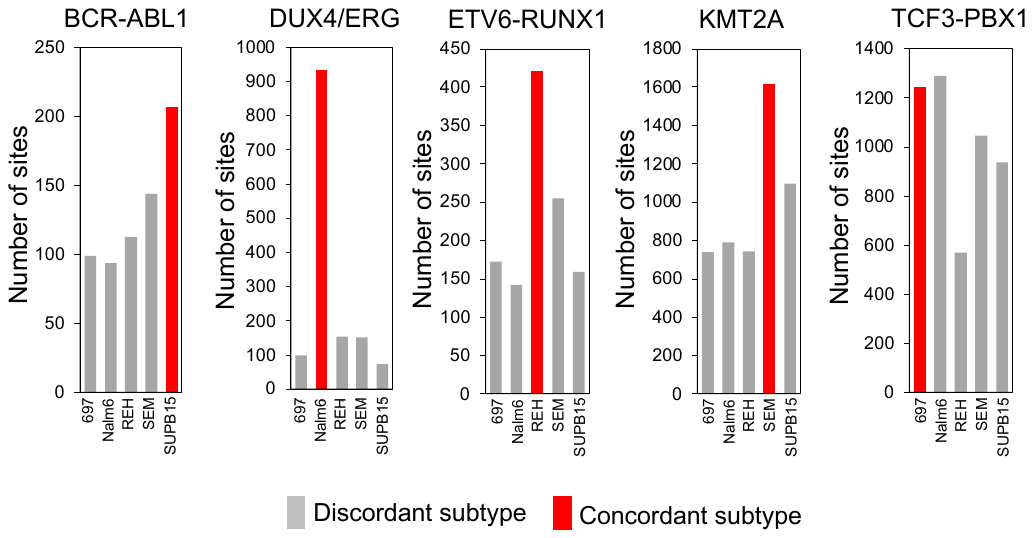

**Figure S3.** Across a panel of B-ALL cell lines (697, Nalm6, REH, SEM and SUPB15; x-axis), bar plots delineate the number of (from left to right) BCR-ABL1, DUX4/ERG, ETV6-RUNX1, KMT2A-rearranged and TCF3-PBX1 DAS that exhibit the strongest accessibility in each B-ALL cell line. Red denotes the corresponding B-ALL cell line for each subtype-enriched DAS and gray denotes B-ALL cell lines from opposing subtypes.

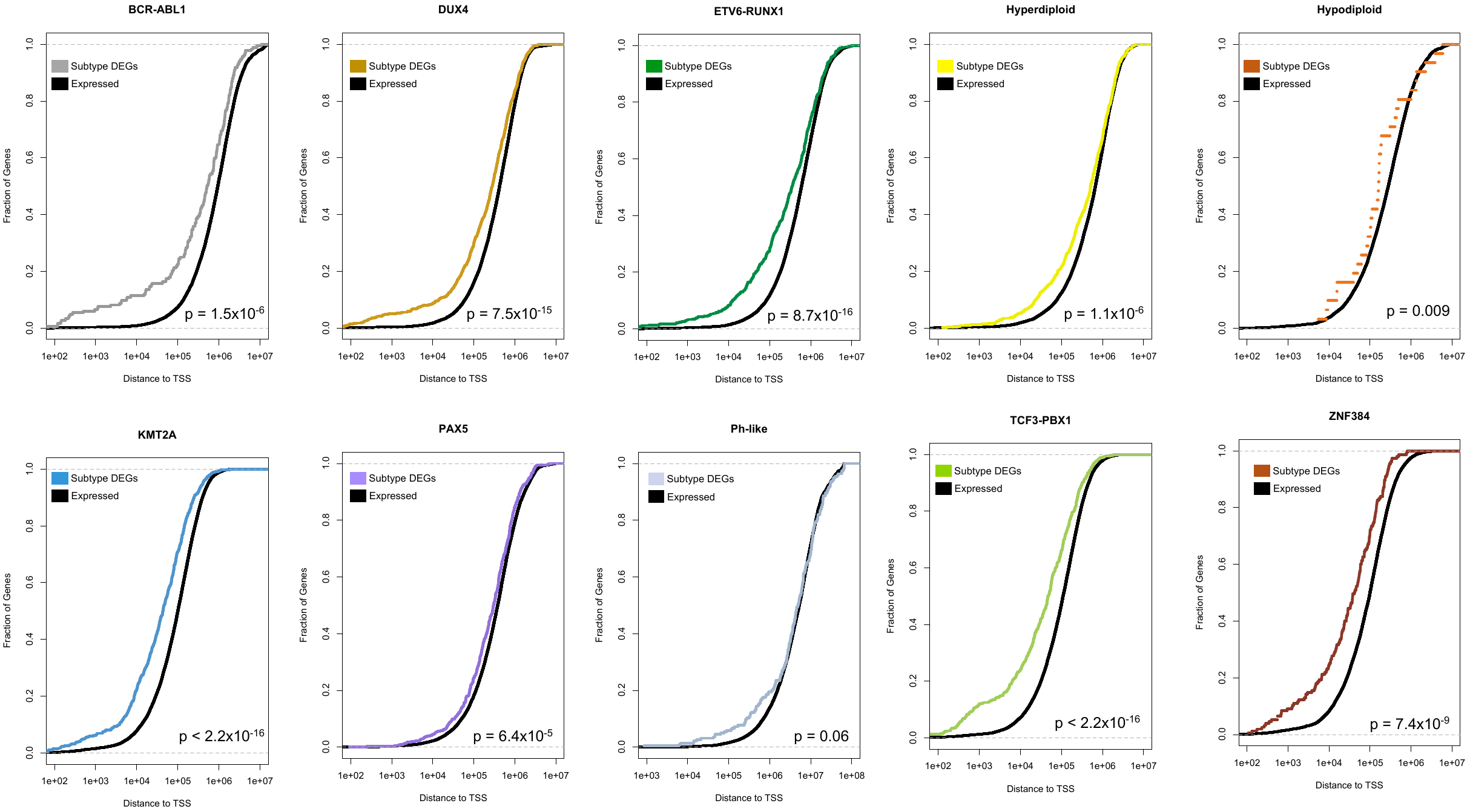

**Figure S4.** Cumulative distribution function comparing the fraction (y-axis) of subtype up-regulated genes (Subtype DEGs; red) and all expressed subtype gene (Expressed genes; black) at different distance cutoffs from subtype-enriched DAS and their transcription start sites (x-axis). Kolmogorov-Smirnov (K-S) p-values are provided on each figure and data is provided for BCR-ABL1, TCF3-PBX1 and ZNF384 ALL.

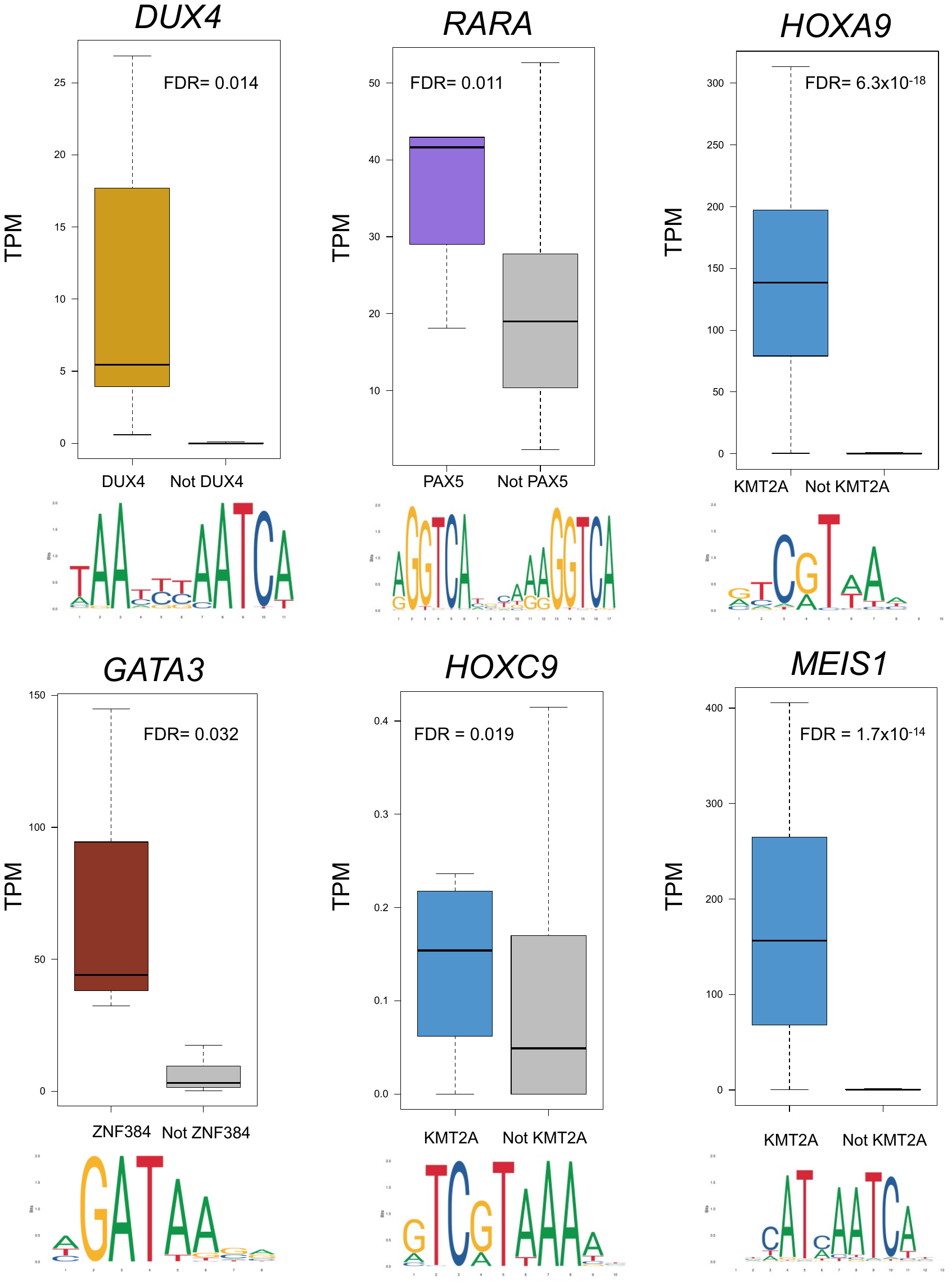

**Figure S5.** RNA-seq transcripts per million (TPM) expression of key TFs with subtype-enriched footprints that are also up-regulated in the corresponding subtype (in red) versus all other subtypes (gray). DESeq2 differentially expressed gene FDR significance values are provided.

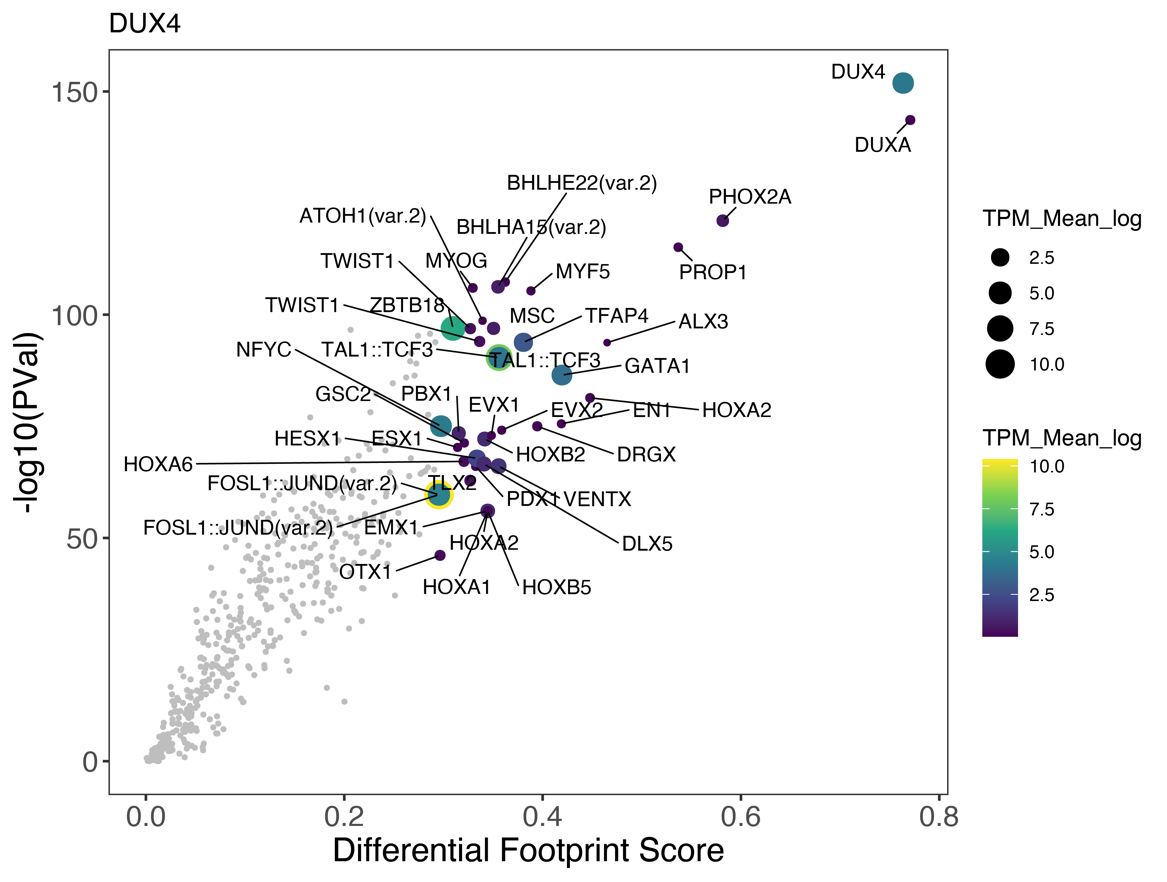

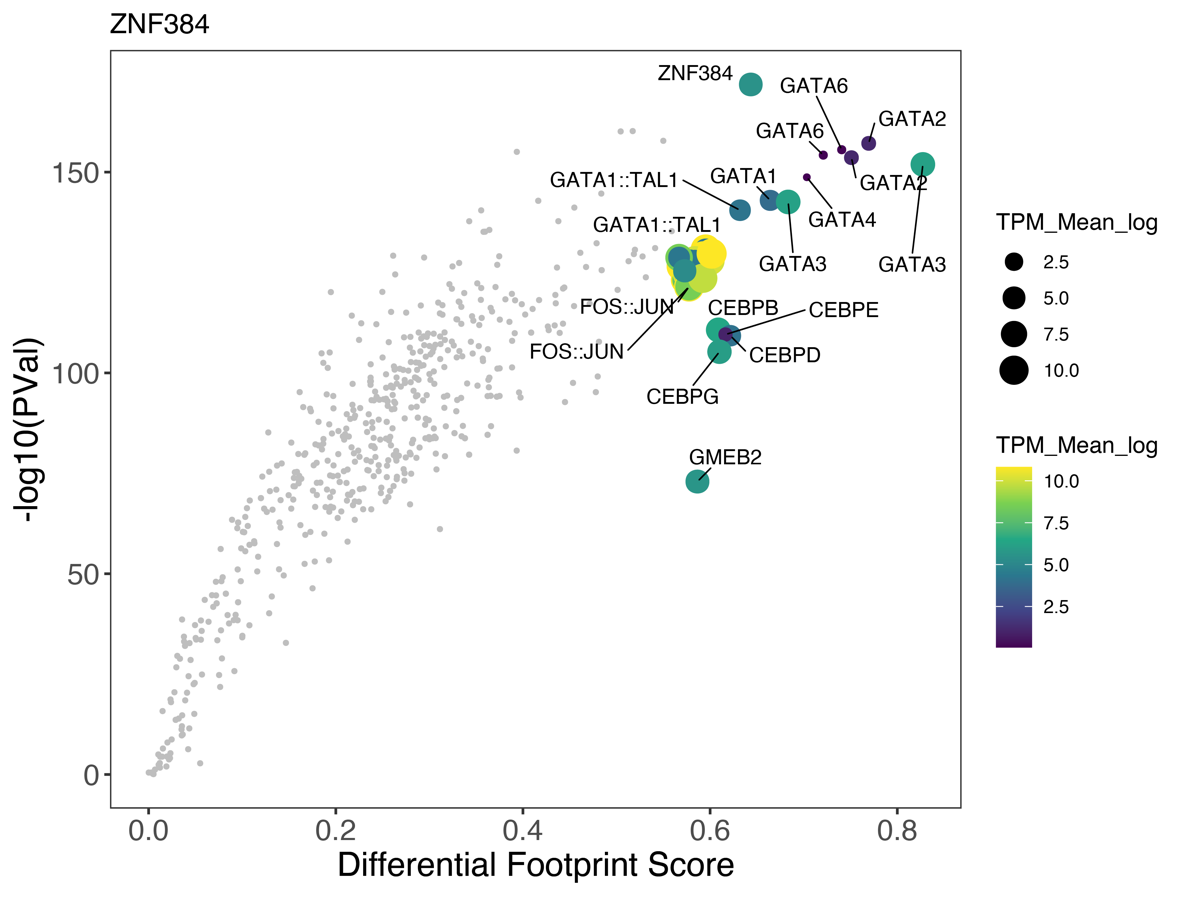

**Figure S6.** Differential footprint score between B-ALL and Pro-B cells is provided on the x-axis and TF footprint significance is provided on the y-axis. TPM transcript abundance of associated TF transcript is shown as both color and size of points. Data is shown for DUX4-enriched DAS (top) and ZNF384-enriched DAS (bottom).

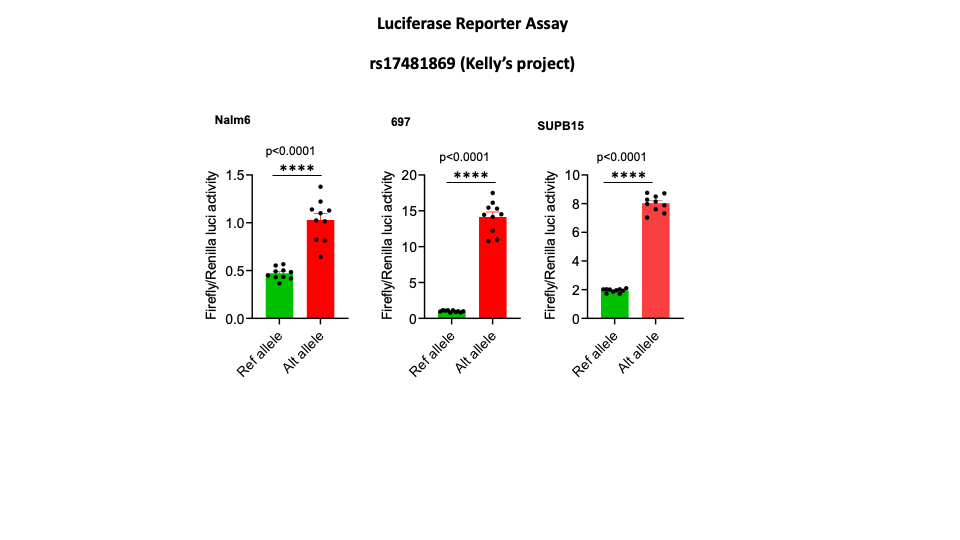

**Figure S7.** Luciferase reporter plasmid activity for SNP rs17481869 comparing reference allele and alternative allele in Nalm6, 697 or SUBP15 B-ALL cell lines. Firefly luciferase signal is normalized to renilla luciferase control.
