## Supplemental Methods for "Epigenomic mapping in B-cell acute lymphoblastic leukemia identifies transcriptional regulators and noncoding variants promoting distinct chromatin architectures"

**Chromatin accessibility landscapes of B-cell acute lymphoblastic leukemia subtypes identifies epigenomic heterogeneity**

Kelly R. Barnett, PhD^1,2^, Robert J. Mobley, PhD^1,2^, Jonathan D. Diedrich, PhD^1,2^, Brennan P. Bergeron, PhD^1,2,3^, Kashi Raj Bhattarai, PhD^1,2^, Wenjian Yang, PhD^1,2^, Kristine R. Crews, PharmD^1,2^, Christopher S. Manring, MBA^4^, Elias Jabbour, MD^5^, Elisabeth Paietta, PhD^6^, Mark R. Litzow, MD^7^, Steven M. Kornblau, MD^5^, Wendy Stock, MD^8^, Hiroto Inaba, MD, PhD^1,9^, Sima Jeha, MD^1,9^, Ching-Hon Pui, MD^1,9^, Charles G. Mullighan, MBBS (Hons), MSc, MD^1,10^, Mary V. Relling, PharmD^1,2^, Jun J. Yang, PhD^1,2,3,11^, William E. Evans, PharmD^1,2^ and Daniel Savic, PhD^1,2,3,11,*^

^1^ Hematological Malignancies Program, St. Jude Children’s Research Hospital, Memphis, TN 38105, USA.

^2^ Department of Pharmacy and Pharmaceutical Sciences, St. Jude Children’s Research Hospital, Memphis, TN 38105, USA.

^3^ Graduate School of Biomedical Sciences, St. Jude Children’s Research Hospital, Memphis, TN 38105, USA.

^4^ Alliance Hematologic Malignancy Biorepository; Clara D. Bloomfield Center for Leukemia Outcomes Research, Columbus, OH 43210, USA

^5^ Department of Leukemia, The University of Texas M. D. Anderson Cancer Center, Houston, TX, USA

^6^ Department of Oncology, Montefiore Medical Center, Bronx, NY 10467, USA.

^7^ Division of Hematology, Department of Medicine, Mayo Clinic, Rochester, MN 55905, USA.

^8^ University of Chicago Comprehensive Cancer Center, Chicago, IL 60637, USA.

^9^ Department of Oncology, St. Jude Children’s Research Hospital, Memphis, TN 38105, USA.

^10^ Department of Pathology, St. Jude Children’s Research Hospital, Memphis, TN 38105, USA.

^11^ Integrated Biomedical Sciences Program, University of Tennessee Health Science Center, Memphis, TN 38105, USA.

*Corresponding author:	Daniel Savic, PhD

Division of Pharmaceutical Sciences

Department of Pharmacy and Pharmaceutical Sciences

St. Jude Children’s Research Hospital

262 Danny Thomas Place

Memphis, TN, 38105

**Data analysis – Regions of Interest Selection**

Accessible chromatin sites analyzed throughout this work were selected using a reproducible ATAC-seq peak summit approach as follows. Peak summits were generated for each subtype on a subtype-merged basis and narrowPeak regions for each individual patient sample. Any subtype-merged peak summit not reproducible among multiple individual patient sample narrowPeaks was excluded for analysis. All reproducible, subtype-merged summits were then extended upstream and downstream to an interval size of 301bp and merged if overlapping. Finally, any interval overlapping hg19 blacklist regions were eliminated yielding the final set of ATAC-seq regions of interest used in further analysis. The ChIPseeker R-package was used for genomic annotation of all genomic intervals throughout this work.

**Data analysis – Pro-B cell and B-ALL Comparison**

DESeq2 using the Wald statistical test was utilized to calculate differential chromatin accessibility between normal Pro-B cells (sourced from GSE122989) B-ALL patient samples. All samples were required to pass an ATAC-seq quality score cutoff to be included in analysis. ATAC-seq quality scores were determined by calculating ATAC-seq read enrichment around transcription start sites Differential chromatin accessibility between normal Pro-B cells and B-ALL patient samples was defined as chromatin regions passing a p-adjusted filter of < 0.05 and an absolute value log2(fold change) >= 2. A variance stabilized transform function within DESeq2 was applied to the ATAC-seq read counts matrix prior to clustering with the pheatmap R-package for visualization.

**Data analysis – Subtype Enriched Chromatin Accessibility**

DESeq2 using the Wald statistical test was utilized to calculate differential chromatin accessibility among B-ALL subtypes. Cohorts representing a single subtype were compared to all other B-ALL patient samples not belonging to the single subtype. This pairwise comparison was completed for all subtypes and samples with sufficient sample numbers (N > 1). Subtype-enriched chromatin accessible regions were required to pass filters of p-adjusted < 0.05 and an absolute value log2(fold change) >= 1. Subtype-enriched regions were additionally required to be exclusively differential in a single subtype, regions appearing as differential in multiple subtypes were excluded. A variance stabilized transform function within DESeq2 was applied to the ATAC-seq read counts matrix specific to subtype-enriched loci prior to visualization with the pheatmap R-package.

**Data analysis – ATAC-QTL Identification**

VCF (Variant Call Frequency) files were sourced from St. Jude Children’s Research Hospital genotyping. Variants in this dataset represented a mixture of both directly genotyped and imputed variants. Imputation was performed via the Michigan imputation server (version 1.6.5) using minimac4 for imputation, eagle-2.4 for phasing and the TOPMed reference panel. The final variant list for analysis with RASQAUL was restricted to variants within B-ALL open chromatin regions yielding 914,406 variant SNPs in total. Allele specific ATAC-seq read counting for open chromatin region SNPs was performed with the RASQUAL supplied helper script which utilizes the GATK ASEReadCounter tool. All SNPs were required to have an imputation quality R^2^ of >= 0.80 for final inclusion after running RASQUAL.

**ATAC-seq**

Amplified DNA libraries were sequenced on an Illumina Nova-seq 6000 where >50M 150bp paired-end reads were obtained for all samples. Adapter trimming was performed using TrimGalore (v0.6.6) with command options “--fastqc --paired”. Read mapping was performed using Bowtie2 (v2.2.9) and hg19 genome index with custom command options “-X 2000 -S”. Read quality filtering was performed with Samtools with the “view” command and options “-q 20 -b”. Sorting was performed with Picard (v1.141) using the “SortSam” command and options “SORT_ORDER=coordinate”. Mitochondrial reads were removed using Samtools “view” command combined with command line filtering, “samtools view -h bam | awk '{if($3 != "chrM"){print $0}}' | samtools view -b - > bam”. Peak calling was performed with MACS2 (v2.1.1 using command and options “macs2 callpeak -t bam -f BAMPE -g hs --nomodel --extsize 200 --SPMR -B”.

**ChIP-seq**

Amplified DNA libraries were sequenced on an Illumina Nova-seq 6000 where >50M 150bp single-end reads were obtained for all samples. Adapter trimming was performed using TrimGalore (v0.6.6) with command options “--fastqc”. Read mapping was performed using Bowtie2 (v2.2.9) and hg19 genome index with custom command options “-X 2000 -S”. Read quality filtering was performed with Samtools with the “view” command and options “-q 20 -b”. Sorting was performed with Picard (v1.141) using the “SortSam” command and options “SORT_ORDER=coordinate”. Mitochondrial reads were removed using Samtools “view” command combined with command line filtering, “samtools view -h bam | awk '{if($3 != "chrM"){print $0}}' | samtools view -b - > bam”. Peak calling was performed with MACS2 (v2.1.1) using command and options “macs2 callpeak -t bam -f BAM -g hs --nomodel --extsize 200 --SPMR -B”.

**Promoter capture Hi-C**

Arima promoter capture HiC (Arima product #s : A510008, A303010, A302010) was performed on B-ALL cell lines (697, Nalm6, RS411, REH, SUPB15, BALL1, SEM) according to the manufacturers provided instructions using unspecified proprietary buffers, solutions, enzymes, and reagents. Briefly, 10 million ALL cells were harvested, suspended in 5ml RT PBS which was brought to 2% formaldehyde by adding 37% methanol-stabilized paraformaldehyde for a 10-minute fixation. The amount of fixed cell suspension equal to 5μg of cell DNA was used for HiC. Cells were lysed with Lysis Buffer and conditioned with Conditioning Solution before their DNA was digested in a cocktail consisting of Buffer A, Enzyme 1, and Enzyme 2. The digested, fixed chromatin was biotinylated using using Buffer B and Enzyme B before being ligated using Buffer C and Enzyme C. The fixed, biotinylated, ligated DNA was then subjected to reversal of crosslinking and digestion of proteins before being purified. 100ul containing 1500ug of purified large proximally ligated DNA was fragmented for 24 cycles (30s on/ 30s off) using a Diagenode Bioruptor Plus bath sonicator. The fragmented DNA was then subjected to two-sided size selection targeting fragments between 200-600bp using AMPure XP DNA purification beads. Size selected DNA was then subjected to biotin enrichment using T1 streptavidin beads. Bead bound, enriched HiC DNA was then subjected to Arima library prep. Briefly, the sample underwent end repair followed by adapter ligation, at which point the sample was then subjected to 10 cycles of PCR amplification. The library DNA was then purified using AMPure XP DNA purification beads. The HiC library was then subjected to Arima promoter capture enrichment. The library was precleared of biotinylated DNA using T1 streptavidin beads before being subjected to promoter enrichment with biotinylated RNA probes. After washing, the captured fragments were then amplified an additional 13 PCR cycles.

**Promoter Capture HiC-seq analysis**

Amplified DNA libraries were sequenced on an Illumina Nova-seq 6000 where >200M 150bp paired-end reads were obtained for all samples. Analysis of promoter capture HiC data was performed using the Arima CHiC pipeline (v1.4, <https://github.com/ArimaGenomics/CHiC>). Briefly, this pipeline uses HiCUP v0.8.0 for mapping and quality assessment of promoter capture HiC data and CHiCAGO to identify significant looping interactions in the promoter capture HiC data using 3kb resolution and adjusted p-value < 0.05. Files were processed at 3kb resolution with command “bash Arima-CHiC-v1.4.sh” with key custom options including: “-W 1 -Y 1 -Z 1 -P 1 -d Digest_hg19_Arima.txt -b human_GW_PC_S3207364_S3207414_hg19.uniq.bed -R hg19_chicago_input_3kb.rmap -B hg19_chicago_input_3kb.baitmap -O hg19”. The following input files: hiccup genome digest (Digest_hg19_Arima.txt), probe design file (human_GW_PC_S3207364_S3207414_hg19.uniq.bed), rmap file (hg19_chicago_input_3kb.rmap), baitmap file (hg19_chicago_input_3kb.baitmap) and corresponding hg19 3kb resolution CHiCAGO design files (*.npb, *.poe, *.nbpb) were sourced from Arima FTP server ( <ftp://ftp-arimagenomics.sdsc.edu/pub/ARIMA_Capture_HiC_Settings/>). All significant intrachromosomal chromatin interactions spanning less than 2Mb were concatenated from all B-ALL cell lines to create a comprehensive library of pan-B-ALL cell line promoter capture chromatin loops. Genomic regions representing separate loop ends were compiled to facilitate overlap determinations with B-ALL patient chromatin accessible regions of interest using “bedtools intersect”.
